# Three new Cenomanian conifers from El Chango (Chiapas, Mexico) offer a snapshot of the geographic mosaic of the Mesozoic conifer decline

**DOI:** 10.1101/2021.09.01.458614

**Authors:** Ixchel González-Ramírez, Sergio R.S. Cevallos-Ferriz, Carl J. Rothfels

**Affiliations:** University Herbarium and Department of Integrative Biology, University of California, Berkeley; Instituto de Geología, Universidad Nacional Autónoma de México

**Keywords:** fossils, Cretaceous, Cenomanian, Cupressaceae, *Dacrydium*, *Microcachrys*, Podocarpaceae, conifer phylogeny, *Sequoiadendron*

## Abstract

**Premise of study:** “El Chango” is a recently discovered quarry that contains extremely well preserved fossils. The Cenomanian age of the locality corresponds to a time when the global flora was transitioning from gymnosperm- to angiosperm-dominated, yet conifers predominate in this locality. These fossils thus provide a rare opportunity to understand the replacement of conifers by angiosperms as the dominant group of plants.

**Methods:** We collected material from El Chango in annual expeditions (2010 to 2014). We selected the three most abundant and best preserved conifer morphotypes and conducted a total-evidence (i.e.,, including molecular and morphological data) phylogenetic analysis of a sample of 72 extant conifer species plus the three fossils. We use these results to inform our taxonomic decisions.

**Results:** We obtained four equally most-parsimonious trees (consistency index = 44.1%, retention index = 78.8%). Despite ambiguous relationships among some extant taxa, the three fossil conifers had the same phylogenetic position in all four most-parsimonious trees; we describe these species as new: *Sequoiadendron helicalancifolium* sp. nov. (Cupressaceae), and *Microcachrys rhomboidea* sp. nov. and *Dacrydium bifoliosus* sp. nov (Podocarpaceae). The ecosystem is interpreted as a coastal humid mixed forest.

**Conclusions:** Our findings contribute to the understanding of Cenomanian equatorial regions, and support the hypothesis of a geographically and ecologically structured “rise of angiosperms”, with conifers remaining dominant in brackish-water and angiosperms becoming dominant in freshwater-ecosystems. These fossils fill in gaps in the evolutionary history of lineages like *Microcachrys*, which we demonstrate occurred in the Northern hemisphere before becoming restricted to its current range (Tasmania).

## INTRODUCTION

Despite being massively outnumbered by extant angiosperms (angiosperms account for ∼ 89% of extant plant species vs ∼ 0.29% for gymnosperms; Crepet and Niklas, 2009; Hassler, 2021) conifers were dominant for the first half of the Mesozoic (Miller, 1977; Taylor et al., 2009). This former ecological dominance is documented in a particularly rich fossil record (e.g., Contreras et al., 2019; Hernández-Castillo et al., 2014; Looy, 2007), which is partially due to the fact that conifers have sturdy structures—i.e., cones, wood, and thick leaves—that are more prone to fossilize than are the more delicate structures of other plant groups. This fossil record reveals that conifers first evolved during the Carboniferous, likely from an ancestor in the Cordaitales, an extinct group of seedplants (Beck, 1988). During the rest of the Paleozoic, conifers increased in diversity and geographic extent, leaving behind traces of lineages that are known only from the fossil record, such as the Walchian and Voltzian conifers (Looy, 2007; Hernández-Castillo et al., 2014). Though most of the Paleozoic conifer lineages went extinct around the Permian-Triassic boundary, the groups that persisted became widespread in the Mesozoic (Beck, 1988; Farjon, 2008). There are records of nine families of conifers in the Mesozoic, including the six extant families (Taylor et al., 2009). During the Triassic and Jurassic, conifers occurred in a wide variety of habitats, including some that are currently dominated by angiosperms. This diversity of habitats was also associated with a diversity of life forms, such as the records of a ruderal herbaceous conifer (Rothwell et al., 2000).

Despite the abundance of conifers in the fossil record, reconstructing their evolutionary history is challenging. First, plant fossils are usually fragmentary and only in very rare cases do different organs (e.g., leaves, stems, roots, reproductive structures) appear connected in the fossil record. This difficulty in determining which pieces go together extends to within-organ assessments—conifers, in particular, can have different types of leaves and leaf arrangement (i.e., phyllotaxy) depending on the position of the branch and whether or not the branch is reproductive (Hernandez, 2006; Farjon, 2008). In addition, even when the whole plant can be reconstructed with some confidence, homology to features of extant groups is often uncertain.

Another challenging aspect about understanding the history of conifers is the temporal and spatial heterogeneity in the availability of fossils (Allison and Bottjer, 2011). For example, areas closer to the equator tend to have thick vegetation that complicates the access to fossil-bearing rock layers. Therefore, the fossil record is biased towards geographic areas where rocks are exposed, like deserts or cliffs, especially those of regions with a long history of paleontological exploration (Europe and North America). In Mexico, for example, most of the plant fossils from the Cretaceous are from northern areas of the country (Villanueva-Amadoz et al., 2014), where deserts dominate the landscape. In this context, the recent discovery of a rich fossil-bearing site in Chiapas (southern Mexico) is particular noteworthy as it increases our latitudinal sample of terrestrial communities in the Cenomanian Therefore, by studying the plant community of El Chango, we can obtain some insight regarding the latitudinal variation of vegetation in the Cenomanian, as well as particular information about the evolutionary history of individual lineages.

This site, named “El Chango”, is Cenomanian in age and contains extremely well preserved fossils of fishes, invertebrates, angiosperms, and gymnosperms (Alvarado-Ortega and Than-Marchese, 2013, 2012; Guerrero-Márquez et al., 2013; Huerta-Vergara et al., 2013; González-Ramírez et al., 2013; Moreno-Bedmar et al., 2014). El Chango’s fossils offer a great opportunity to expand our knowledge about Cenomanian low-latitude floras (Villanueva-Amadoz et al., 2014) filling in an important gap both temporally (i.e., Cenomanian) and geographically (southernmost North America). In this paper we describe three new species of conifers from El Chango and infer their phylogenetic position and taxonomic affinities based on a total-evidence phylogenetic analysis incorporating 72 extant conifer representatives. These new species expand our records of the temporal and geographic occurrence of the lineages they belong to and deepen our understanding of Cenomanian biogeographic patterns. In particular, they provide insight into the identity and distribution of conifers in the middle of the angiosperm rise to dominance.

### Geological framework

El Chango is located in Chiapas, Mexico (16°34.14′N, 93°16.11′W) and belongs to the Cintalapa Formation, in the Sierra Madre Group (Moreno-Bedmar et al., 2014). The sediments that formed the Sierra Madre Group were deposited during the Albian-Santonian, in the mid-Cretaceous, ∼113 to 83.6 Mya (Steele and Waite, 1985). The rocks of El Chango specifically are Cenomanian in age (100.5–93.9 Mya) based on the presence of two stratigraphic markers: the ammonites *Graysonites* and *Metengonoceras* (Moreno-Bedmar et al., 2014). The known section of El Chango is 54 m thick and it is composed primarily of marine laminated dolomites with sporadic flint levels (Moreno-Bedmar et al., 2014). The depositional environment has been interpreted as a coastal lagoon with ephemeral freshwater influx (Vega et al., 2006). El Chango and adjacent quarries like “El Espinal” are considered *Konservat Lagerstätten* due to the excellent preservation of their fossils (Díaz-Cruz et al., 2016). In these layers, scientists have found fishes with soft tissue preservation (Díaz-Cruz et al., 2016; Alvarado-Ortega and Than-Marchese, 2012, 2013; Vega et al., 2003) and many different types of fossil plants, including a variety of angiosperms similar to Arecaeae, Bignoniaceae, Combretaceae, Myrtaceae, and seagrasses in Cimodoceaceae and Hydrocharitaceae (Guerrero-Márquez et al., 2013). However, conifers are the most common group of plants present (González-Ramírez et al., 2013).

## MATERIAL AND METHODS

### Specimen collection and curation

The members of the paleobotany lab of Instituto de Geología, UNAM, and UNICACH (Universidad de ciencias y artes de Chiapas) obtained fossil samples from El Chango in field expeditions from 2010 to 2014. We focused on sampling the 27–30 and 38–41 meter horizons of the stratigraphic column published by Moreno-Bedmar et al. (2014), which are known to contain many plant fossils. We selected three conifer morphotypes based on their abundance and completeness as the subject of this work. We observed and photographed the specimens using a Zeiss Stemi DV4 and Olympus SZH microscope. When cuticles were preserved, we eliminated the rock matrix using HCl and HNO_3_ to isolate the cuticles, following the method developed by Porras Carrasco (2012). The fossil specimens were deposited in the Colección Nacional de Paleontología at the Instituto de Geología, UNAM.

### Phylogenetic analysis

We performed a phylogenetic analysis based on morphological and molecular data from the three fossil morphotypes, 72 extant species of conifers, and one outgroup (*Ginkgo biloba* L.) to investigate the phylogenetic position of the fossils, and to inform our taxonomic treatment (Appendix 1). The 72 extant taxa represent the morphological and phylogenetic diversity of living conifers, with a particularly dense sampling of Cupressaceae and Podocarpaceae because of the morphological similarity of the fossils with members of these families.

### Data matrix assembly

Our molecular data consist of the plastid *matK* and *rbcL* regions, obtained from Genbank (see accession numbers in the Supplemental Material). We aligned the markers using MAFFT v7.409 (Katoh et al., 2002), and adjusted the alignment manually with AliView v.1.26 (Larsson, 2014); ambiguous areas of the alignment were excluded from subsequent analyses. For the *matK* alignment, unambiguous indels were recoded manually following the simple gap recoding approach of Simmons and Ochoterena (2000). For fossil terminals, molecular characters were coded as unknown (“?”).

Our morphological data consist of fifty-seven categorical characters and nine continuous characters. We defined the categorical characters and their states based on previous works (e.g., Hart, 1987; Gadek et al., 2000; Little et al., 2004; Farjon, 2005). Nevertheless, because some of these works focused on single conifer families, we redefined some character states to accommodate comparison among families. A complete list of characters and character states used in this study can be found in the Supplemental Material 2. The nine continuous characters were: (1) length of ovulate strobilus; (2) width of ovulate strobilus; (3) seed length; (4) seed width; (5) length of pollen strobilus; (6) width of pollen strobilus; (7) length of mature leaves; (8) width of mature leaves; and (9) number of ovuliferous complexes (i.e., ovuliferous scale + bract taken as a unit). For each continuous character, we obtained 10 measurements from herbarium samples to estimate the mean and variance in that character for each taxon. For the fossils—especially for the cone characters—we often had to rely on one to three measurements. These continuous data were treated following the methodology proposed by Goloboff et al. (2006), assigning ranges of continuous traits (mean ± sd) to each terminal. We assigned character states for all the extant species based on examination of herbarium specimens from MEXU (National Herbarium of Mexico) and NYBG (Herbarium of the New York Botanical Garden), and a specialized literature search. The final data matrix is available in two repositories: in Github (https://github.com/ixchelgzlzr/coniferas_el_chango and Dryad (XXXX).

### Analyses

We performed a maximum-parsimony analysis in TNT (Goloboff et al., 2005). We used 1000 different starting trees and applied “new technology search algorithms” as follows: ratchet (10 repetitions), sectorial search, drift (10 repetitions), and tree fusing (three rounds). We specified all the characters as equally weighted and unordered except for characters 3, 16, 36, and 44, which we treated as ordered because we assume that they evolve in a sequence. We inferred 100 bootstrap trees, also with TNT. We calculated bootstrap support values in R (R Core Team, 2020) using the package phangorn (Schliep et al., 2017). We also edited TNT output in R (Wickham, 2019; Revell, 2012) to obtain a tree in Nexus format, and visualized it in FigTree (Rambaut, 2014). The R script is available in Github (https://github.com/ixchelgzlzr/coniferas_el_chango)

## PHYLOGENETIC RESULTS

The *rbcL* and *matK* alignments consisted of 1280 and 1477 positions, respectively. From those, 278 and 713 sites, respectively, were parsimony-informative. We recoded 19 indels, 18 of which were parsimony-informative. The total number of parsimony-informative characters in the molecular data set was 1009. The morphological data consisted of 57 discrete and nine continuous characters, all of which were parsimony-informative.

When we analyzed the concatenated matrix (molecular + morphological data) for the 75 terminals (72 extant species + three fossils) we obtained four equally most-parsimonious trees (MPTs). These trees had a total length of 3962.398 steps, a consistency index (CI) of 44.1% and a retention index (RI) of 78.7% (see Figure 1). The four MPTs differed in the position of *Austrocedurs chilensis* and the position of *Cupressus sempervirens + Cupressus funebris* with respect to the *Callitropsis* clade. All four MPTs resolve the currently recognized extant conifer families (i.e., Araucariaceae, Cupressaceae, Pinaceae, Podocarpaceae, Sciadopytiaceae, and Taxaceae) as monophyletic (Figure 1).

**Figure 1:**
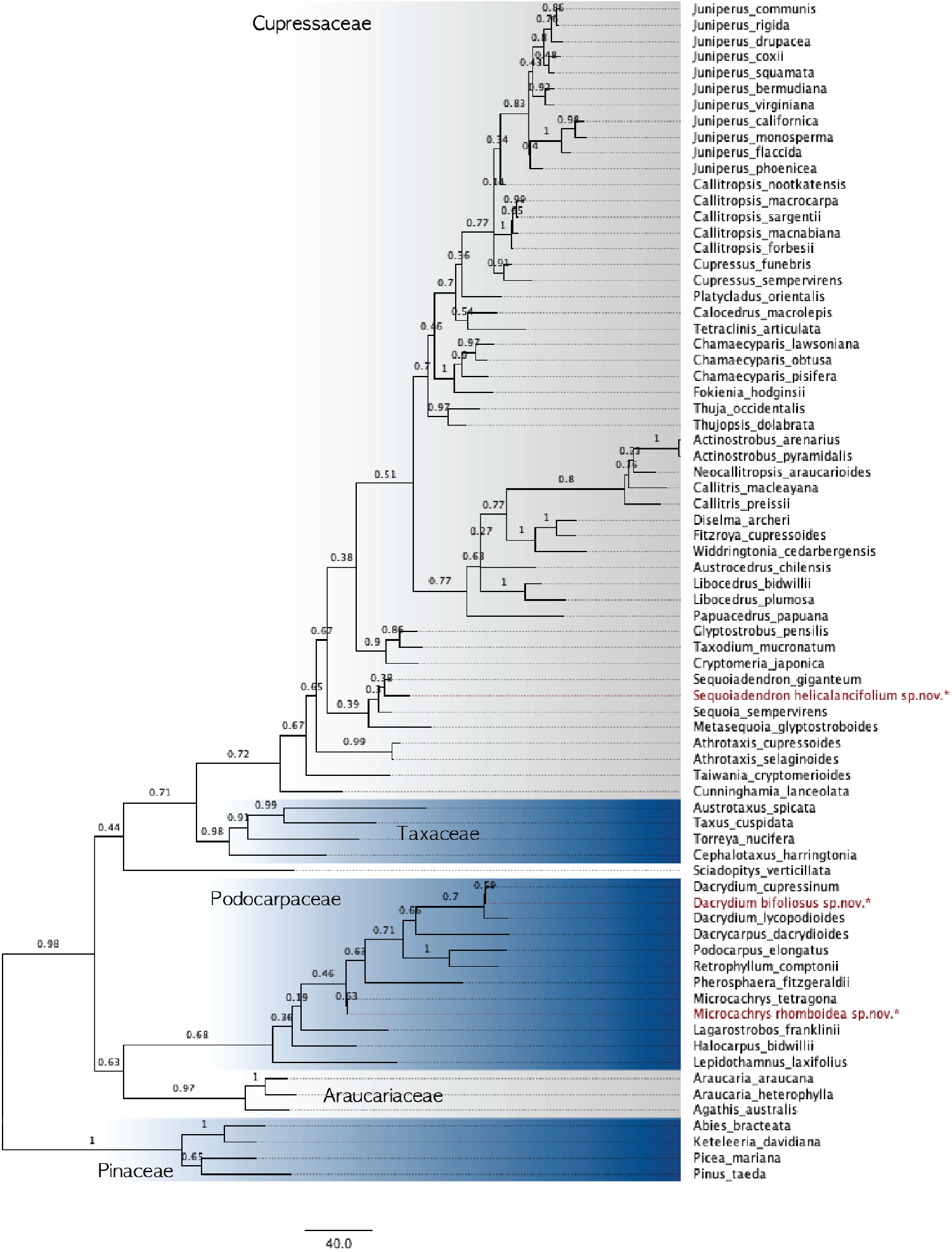
Phylogenetic relationships of 72 extant and three fossil conifers (in red). The relationships were inferred from a parsimony analysis of a combined morphological + molecular dataset. The branch lengths are proportional to the number of steps and the bootstrap values are shown above each branch. The tree was rooted with *Ginkgo biloba*, which was subsequently pruned. The different colors highlight the different families of conifers.

Despite the ambiguous relationships of some of the extant taxa, the three fossils had the same phylogenetic position in all four MPTs. The bootstrap values of the branches uniting the fossils with their sister groups ranged from 0.30–0.59. These bootstrap values are low for standard molecular phylogenetic inferences, but considering the amount of missing data in the fossils (i.e., the complete lack of molecular information and extensive missing morphological data), we consider the consistent placement of the fossils in the MPTs as strong evidence of their taxonomic affinities, which we describe below.

### Taxonomic descriptions

> Kingdom **Plantae** Jussieu (1774)
>
> Division **Tracheophyta** Sinnot (1935)
>
> Subdivision **Spermatophytina** Cavalier-Smith (1998)
>
> Class **Pinopsida** Burnett (1835)
>
> Order **Araucariales** Gorozh. (1904)
>
> Family **Podocarpaceae** Endl. (1847), nom. cons.
>
> Genus ***Microcachrys*** Hook. f. (1845)
>
> Species ***Microcachrys rhomboidea*** González-Ramírez sp. nov.
>
> Holotype: Provisional number M2-007. Figure 2
>
> Paratypes: Provisional numbers M2-002, M2-003, M2-006, M2-019, M2-023,

**Figure 2:**
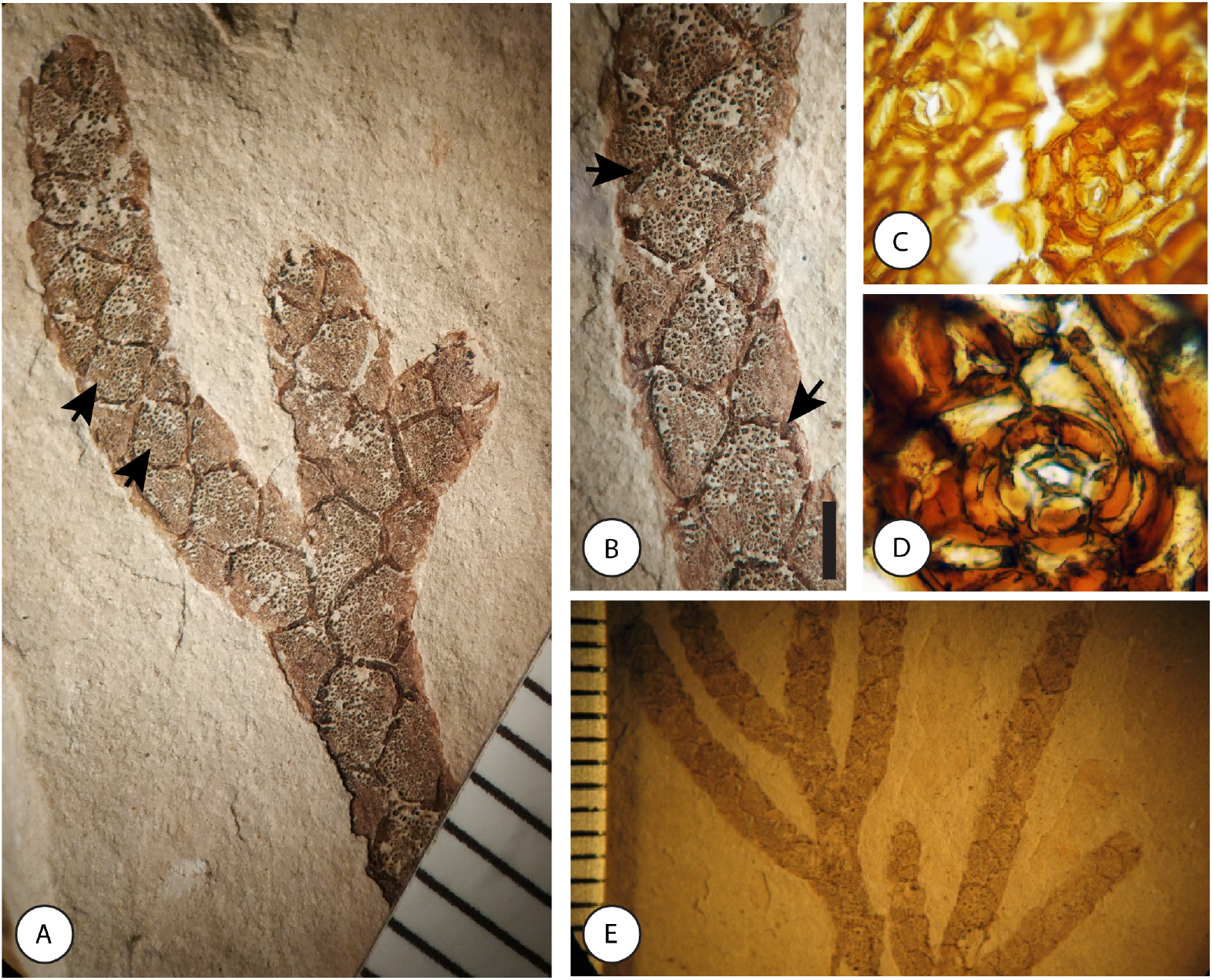
Microcachrys rhomboidea. Provisional number M2-007. A. Branch with leaves. The arrows point to the typical rhomboid like pattern formed by leaf phyllotaxy. B. Magnification of the specimen showed in A (scale bar = 1mm). Arrows point to the cuticle projections on the tip of the leaves that show as darker bands in the fossils. C(20X) and D (40X) show a segment of isolated cuticle. The epidermal cells are rectangular, and the stomata are rounded with two cycles of cells. E. Sample illustrating the regularly spaced branching. The rulers are in millimeters.

#### Locality

México, Chiapas, Municipio de Ocozocuautla, El Chango Quarry (16°34’14” N, 93°16’11” W). Locality number: 3923.

#### Stratigraphic occurrence and age

Laminated limestones of the upper section of Cintalapa Formation, Sierra Madre Group (*sensu* Moreno-Bedmar et al., 2014). Cenomanian, between 100.5 and 97.2 Mya.

#### Diagnosis

Scale-like leaves in an opposite criss-crossed arrangement, decurrent and adpressed. Leaves with a distal projection as in *Microcachrys tetragona* (Hook.) Hook. f. Differs from *M. tetragona* in leaves slightly longer and wider. Also, the arrangement and very regular spacing of leaves produce a characteristic pattern (a line of diamonds) on twigs (arrows on figure 2A).

#### Description

Ultimate-order branches alternate and regularly spaced along the penultimate-order branches (Figure 2A). Leaves simple, sessile, and in opposite pairs, slightly imbricate and completely appressed to the branches (as opposed to having the leaf fully or partially free). The imbrication and very regular spacing of the leaves produces a rhomboid pattern along the branch that is distinctive of this species (Fig.2A. Scale-like leaves, as wide as long (∼2mm). Leaf apex obtuse with usually a rounded tip. Leaf margins smooth with a continuous distal band—darker than the rest of the fossil—that is interpreted as a cuticle projection (Fig.2B). This feature is best observed in fully grown leaves, as opposed to the young leaves that occur in the distal part of the branches.

Leaves without noticeable variation in shape: no dimorphism between leaves in facial versus lateral position, and no differences in leaves belonging to different branch orders. Resin glands absent. Stomata dispersed and rounded; oclusive cells surrounded by two cycles of irregularly shaped cells, the first cycle of which is usually formed by six cells and the second cycle by nine to 11 cells. Morphology of the stomata is consistent with the cyclocytic type (Porras Carrasco, 2012). Epidermal cells rectangular, with smooth walls, and arranged in uniform rows. Reproductive structures unknown.

#### Etymology

The specific epithet *rhomboidea* refers to the rhomboid pattern that the leaf arrangement produces.

#### Notes

We resolve this species as the sister group of *Microcachrys tetragona* (Figure 1, Bootstrap support = 0.53). Furthermore, the characters of the fossils are consistent with the circumscription of the genus *Microcachrys* (Eckenwalder, 2009). However, the reproductive structures of this fossil morphotype are unknown and its placement in *Microcachrys* should be re-evaluated with the discovery of more fossils.

> Genus ***Dacrydium*** Lamb. (1807)
>
> Species ***Dacrydium bifoliosus*** González-Ramírez sp. nov.
>
> Holotype: Provisional number M1-003. Fig. 3A
>
> Paratypes: Provisional numbers M1-001 (Fig. 3B), M1-002 (Fig. 3E), M1-004, M1-005

**Figure 3:**
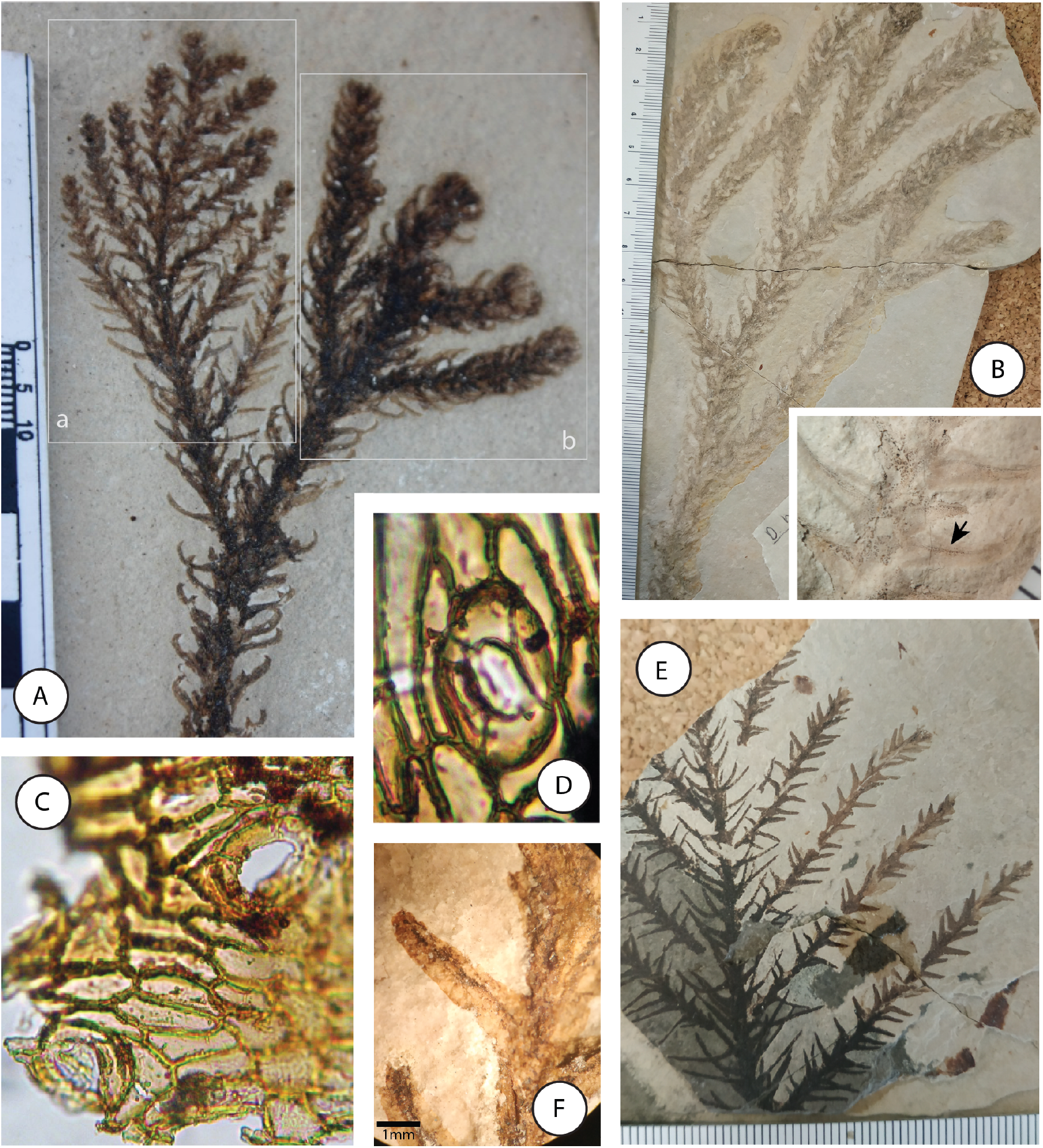
Dacrydium bifoliosus. A. Provisional number: M1-003. Fossil showing the two different branch morphologies (a and b) in organic connection. B. Provisional number: M1-001. Shoot morphology “b”, the most common among the fossils of this locality, with a zoom image where the mid-vein can be observed (arrow). C and D illustrate the cuticles of this morphotype at 20X and 40X respectively. E and F (provisional number: M1-002) show the shoot morphology “a” where branches are regularly arranged in a single axis and leaves are straight.

#### Locality

México, Chiapas, Municipio de Ocozocuautla, El Chango Quarry (16°34’14” N, 93°16’11”O). Locality number: 3923.

#### Stratigraphic occurrence and age

Laminated limestones of the upper section of the Cintalapa Formation, Sierra Madre Group (*sensu* Moreno-Bedmar et al., 2014). Cenomanian, between 100.5 and 97.2 Mya.

#### Specific diagnosis

Ultimate and penultimate branches similar to *Dacrydium lycopodioides* Brongn. & Gris. Differs from that species in that leaves of penultimate branches are incurved. Ultimate branches flattened with shorter distal branches, thus creating an arrow-shaped outline to the branching system. Ultimate branches flattened and very regularly spaced.

#### Description

Branches and leaves dimorphic. In penultimate and higher order branches, the branching pattern is irregular and occurs in different planes. The leaves of these branches are simple, sessile, awl-like, falcate, usually more than 8mm long, helically attached to the branch (forming an acute angle between the leaf axis and the branch), not imbricate, and slightly adpressed. The base of these leaves is approximately twice as wide as the distal portion; the apex is acute and usually pointy, but sometimes rounded. These leaves have a single central vein, smooth margin, and there is no evidence of resin glands.

In ultimate and sometimes penultimate branch orders the branching pattern is almost opposite, regularly spaced, and occurring in a plane. The leaves of these branches are simple, sessile, opposite, regularly spread in a single plane, and slightly appressed to the branch, forming an acute angle. These leaves are awl-like, non-falcate, usually less than 5 mm long and similar width at the distal and the proximal ends; the apex acute and pointy. These leaves have a single mid-vein, smooth margin, and no evidence of resin glands.

Stomata dispersed and ovate. Occlusive cells surrounded by one or two cycles of elongated cells, with typically four cells in the first cycle and between six and eight cells in the second cycle. The morphology of the stomata is consistent with a cyclocytic type. The epidermal cells are rectangular with smooth walls and arranged in regular rows.

The reproductive structures of this species are unknown.

#### Notes

This fossil species’ circumscription was challenging. During the first two years of fieldwork, we collected only fragmented specimens, and from this material we distinguished two forms (Figure 3B and E). The two forms were most clearly distinct in their branch arrangement (in one plane and shortening towards the tip, forming an arrow versus irregular and three-dimensionally branching) and leaf shape (linear versus falcate). In subsequent years it became clear that this original distinction was blurry as we found fossils with intermediate traits, and ultimately we discovered individual fossils (Figure 3) that showed both types of branches. The last piece of evidence that the two forms represented the same species came from the cuticles: both types of branches have the same cuticular traits (Figure 3C and D). The phenomenon of leaf- and branch-dimorphism is common in conifers and is associated with the age, season, and reproductive status of the branches (Hernandez, 2006; Farjon, 2010; Eckenwalder, 2009). The placement of this fossil in the Podocarpaceae is based on the phylogenetic analysis (Figure 1), where this fossil was resolved within *Dacrydium* based on the the uniform length of the branching pattern (character 6) and the epidermal cell arrangement (character 24) and as the sister group of *Dacrydium cupressinum* Sol. ex G. Forst. based on leaf shape (character 10). Furthermore, the known traits of this morphotype are consistent with the traits of the members of *Dacrydium*, and the extant species of this genus present leaf and branch differentiation similar to that observed in the fossils (Eckenwalder, 2009). While we resolve this species within Podocarpaceae, it is important to recognize that it is also similar to species in Cupressaceae, such as *Cryptomeria japonica* (Thunb. ex L. f.) D. Don and *Glyptostrobus pensilis* (Staunton ex D. Don) K. Koch. Once again, it is important to continue fieldwork in this locality, as finding reproductive structures associated with this morphotype would provide decisive evidence to support or reject its current placement.

> Order **Cupressales** Link (1829)
>
> Family **Cupressaceae** Gray (1822)
>
> Genus **Sequoiadendron** J. T. Buchholz, nom. cons. (1939)
>
> Species ***Sequoiadendron helicalancifolium*** González-Ramírez sp. nov.
>
> Holotype: Provisional number M3-001 Fig. 4D and Fig. 4E
>
> Paratypes: Provisional numbers M3-002 (Fig. 4A), M3-003 (Fig. 4B), M3-004 (Fig. 4C),

**Figure 4:**
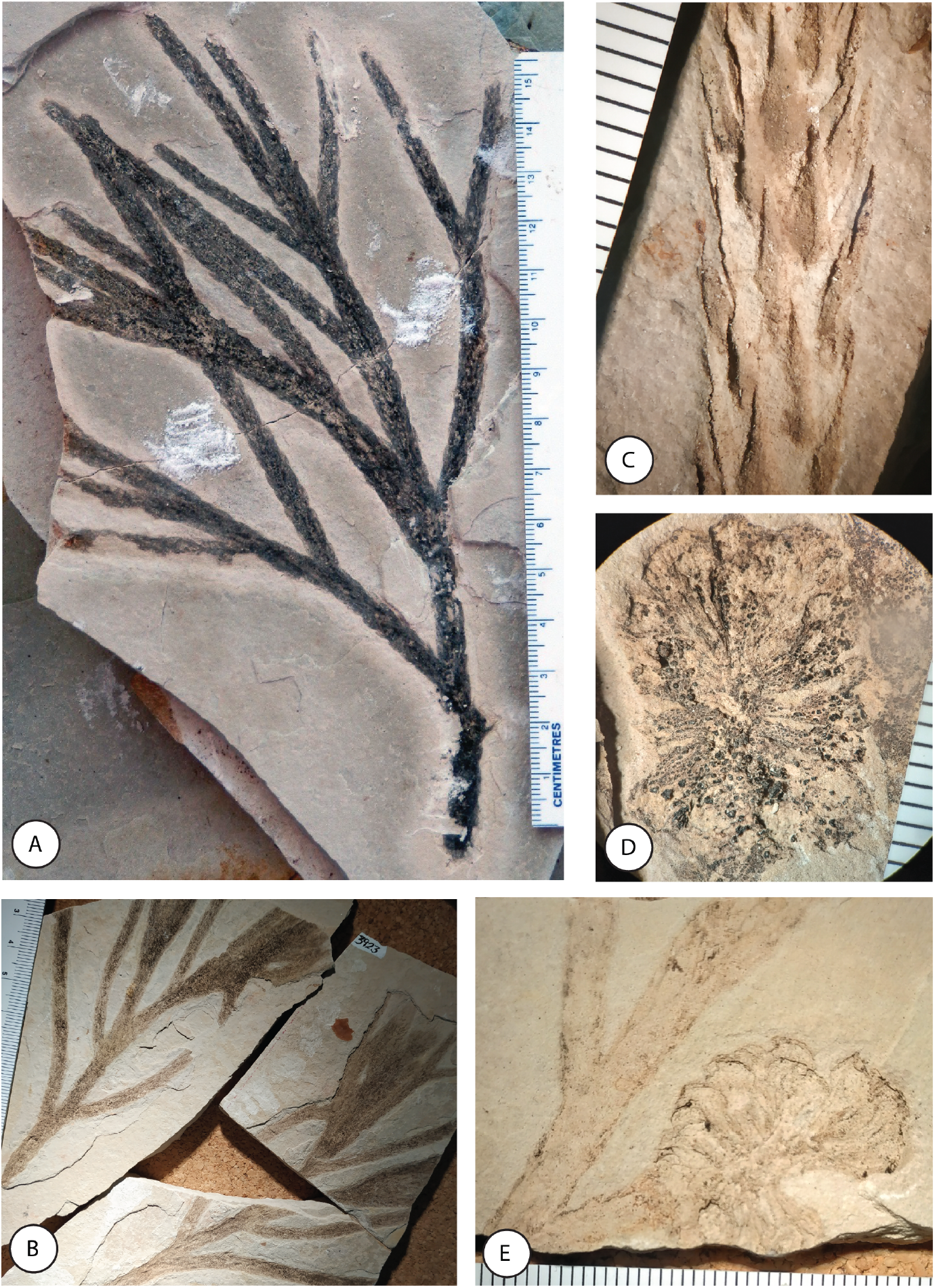
Sequoiadendron helicalancifolium. A (provisional number M3-002) and B (provisional number M3-003). Foliage with dichotomous branching. C. Close-up to the awl-like leaves (provisional number M3-004). D. Ovuliferous cone where the peltate ovuliferous complexes can be observed (provisional number M3-001). E. Ovuliferous cones in organic connection with the foliage (provisional number M3-001).

#### Locality

México, Chiapas, Municipio de Ocozocuautla, El Chango Quarry (16°34’14” N, 93°16’11”O). Locality number: 3923.

#### Stratigraphic occurrence and age

Laminated limestones of the upper section of the Cintalapa Formation, Sierra Madre Group (*sensu* Moreno-Bedmar et al. (2014). Cenomanian, between 100.5 and 97.2 Mya.

#### Diagnosis

Awl-like leaves with helicoidal arrangement, as in *Sequoiadendron giganteum* (Lindl.) J. Buchholz. Leaves differ from *S. giganteum* in being larger (mean=8.2mm) and having a more pointed apex. Also, leaves are generally more imbricate than in *S. giganteum*—they overlap for almost half the leaf length. In addition, the ovulate strobilus is smaller (31mm longer and 23mm wide) than in *S. giganteum*, and globose.

#### Description

The phyllotaxy and dichotomous branch arrangement is similar across the different branch orders (i.e., no branch dimorphism was observed). Leaves simple and sessile, with helicoidal arrangement. Leaves imbricate (covering up to a half of the supra-adjacent leaves), and adpressed approximately half the length of the leaf, but with free tips. Leaves awl-like, longer (mean = 8.2 mm) than wide (mean = 1.6 mm). The apex of the leaves is acute and angular. Leaf margin entire. Resin glands absent. Cuticle-associated traits unknown. Ovulate cone terminal in a lateral branch, globose, dehiscent, 16.9 mm long and 14.5 mm wide (cone measurements from one specimen). No clear differentiation between bract and scale, thus each appendage is referred to as ovulate complex. Ovulate complex peltate, 16–19 in number, helically arranged on a central axis; the axis of each ovulate complex widens towards the distal part.

#### Etymology

The specific epithet describes the shape of the leaves—awl-like and helically arranged—which is one of the most characteristic traits of this species.

#### Notes

The leaves of this fossil have a shape, arrangement, and position similar to *Sequoiadendron giganteum* and they differ in that the fossil leaves are slightly longer and more imbricate. The ovulate cone of *S. helicalancifolium* is similar to the ovulate cone of extant Cupressaceae species in that it has peltate ovuliferous complexes. In particular, it is similar to the extant *Athrotaxis*, *Fokienia*, *Sequoia*, and *Sequoiadendron*. The distal shield of the ovuliferous complex of the fossil is square to rectangular, consistent with *Sequoia* and *Sequoiadendron*. The phylogenetic analysis supports a relationship between the fossil and the sequoioid conifer clade, placing it as the sister group of *Sequoiadendron giganteum*. Given this phylogenetic evidence and the morphological congruence, we name this fossil as a new species of the genus *Sequoiadendron*.

## DISCUSSION

### Implications of the fossil affinities

The three fossils described in this paper closely resemble extant groups of conifers, supporting our current understanding that the traits that distinguish the modern conifers—as opposed to the Paleozoic Voltzian conifers—evolved early in the Mesozoic (Taylor et al., 2009). Furthermore, our findings support the current classification of conifers in two orders: Voltziales—including the Paleozoic conifers—and Coniferales—including the Mesozoic and Cenozoic species (Taylor et al., 2009). Similarly, the fact that the three fossil conifers included in this analysis are sister groups of different extant clades demonstrates that at least some modern genera were clearly differentiated by the Cenomanian. Indeed, other authors (e.g., Leslie et al., 2012) have suggested that the evolution of conifer genera was driven by the break up of Pangaea that occurred ca. 60My before the Cenomanian, in the Jurassic.

The co-occurrence of the three lineages of conifers we describe for El Chango might seem surprising at a first glance, given that extant members of *Microcachrys* and *Dacrydium* occur mainly on islands like New Caledonia, New Zealand, Borneo, and Tasmania, while *Sequoiadendron* is restricted to the Sierra Nevada range in California. Nevertheless, there is substantial evidence that both sequoioid and podocarpoid conifers were much more widespread in the past. In particular, our discovery of *M. rhomboidea* constitutes the oldest macrofossil of *Microcachrys* and fills a gap on the macrofossil record of a lineage that has an broad pollen record that extends geographically to the Cenozoic of Alaska (Reinink-Smith and Leopold, 2005) and temporally to the Jurassic (Truswell and Macphail, 2009), but for which the oldest previously reported macrofossil was from the Cenozoic (Carpenter et al., 2011). Our results are also congruent with divergence-time estimates that place the origin of *Microcachrys* in the Mesozoic (Turner and Cernusak, 2011).

### El Chango flora in a global context

Our current understanding is that the late Cretaceous (Cenomanian to Masstrichtian) was the period of Earth history when angiosperms became the dominant group of terrestrial vegetation. Therefore, it is particularly important to study Cenomanian floras to understand the tempo, mode, and specific drivers of this global transition. One outstanding pattern of the fossil assemblage found in El Chango is that conifers outnumber angiosperms by approximately five-to-one. Determining whether this pattern is the result of taphonomic bias would require a detailed taphonomic study, but it seems safe to conclude that conifers were an important—if not dominant—element of the canopies of El Chango. This abundance of conifers contrasts with other reported Cenomanian floras—for example in Argentina (Iglesias et al., 2007), Utah (Rushforth, 1971), and Russia Moiseeva (2010)—where angiosperms were the dominant group.

Nevertheless, our findings are consistent with the idea that the “rise” of angiosperms was geographically structured, a pattern that Coiffard et al. (2006, 2012) observed in mid-Cretaceous European fossil assemblages where gymnosperms dominated coastal environments while angiosperms were prominent in freshwater environments. Our interpretation of El Chango as a conifer-dominated coastal forest with potentially swampy species like *Microcachrys rhomboidea* supports the idea of conifers remaining dominant in coastal ecosystems during the Cenomanian.

### A glimpse of the Cenomanian forests of El Chango

Although we will never have a full picture of the El Chango ecosystem, our studies provide some insights about the organisms that lived there and the environmental conditions they likely experienced (Fig. 5 is a reconstruction of the terrestrial landscape associated with the El Chango deposits). First, the co-occurrence of marine and terrestrial fossils is strong evidence that the fossil community of El Chango constitutes a death assemblage, a “tanatocenocis” (i.e., at least some of the organisms were transported before deposition and fossilization). The type of sediments is consistent with a coastal lagoon environment (Alvarado-Ortega and Than-Marchese, 2013, 2012; Alvarado-Ortega et al., 2009) and the degree of completeness and the abundance of the plant fossils suggest that the plant remains were transported from a nearby location by moderately calm waters. So, the plants that we find in El Chango probably lived near their deposition site in a coastal forest.

**Figure 5:**
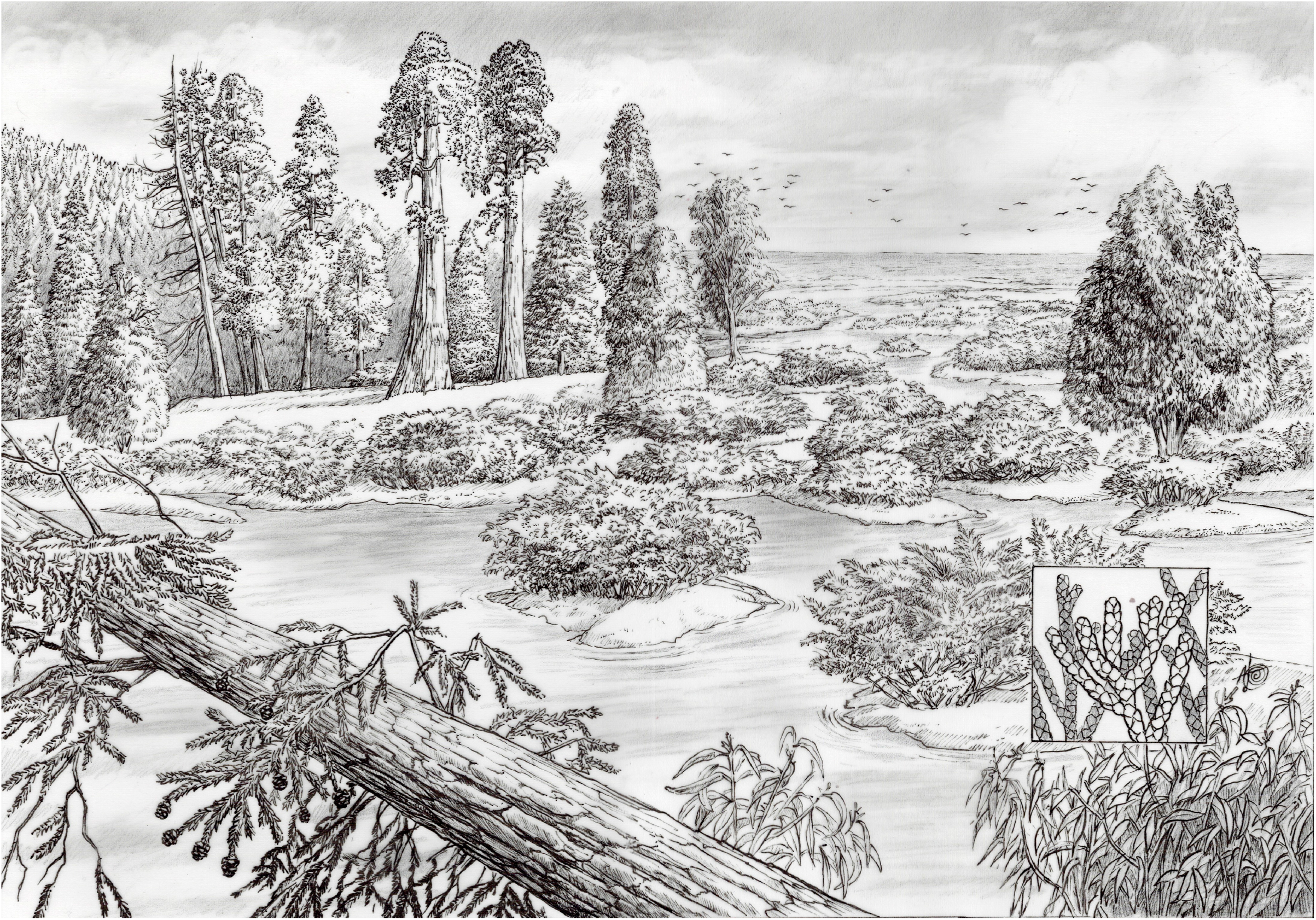
Reconstruction of El Chango site in the Cenomanian. *Sequoiadendron helicalancifolium* sp. nov. (top left) is represented as a tall tree similar to extant redwoods and sequoias. A close-up of its foliage can be seen in the fallen branch on the bottom left. *Dacrydium bifoliosus* sp. nov. appears as a shorter tree (middle right). And *Microcachrys rhomboidea* sp. nov. is illustrated as a small shrub (central part of the image) and a close-up of its branches is on the bottom right. The plants live in a coastal environment associated with a lagoon. Drawing by Aldo Dominguez.

We know, based on estimates from oxygen isotopes, that the surface-temperature of the ocean adjacent to Chiapas during the Cenomanian was warmer than today—around 31 to 36°C (Clarke and Jenkyns, 1999; Laugié et al., 2020), likely because this region was in the path of warm water currents incoming from the Equatorial Atlantic Ocean (Laugié et al., 2020)—so the El Chango forest probably grew in an environment with mild temperatures. The living relatives of the fossils—e.g., *Sequoia sempervirens*, *Sequoiadendron giganteum*, *Dacrydium dacrydioides*, and *Microcachrys tetragona*—are tolerant of a wide range of temperatures but dependent on moist environments. Therefore, the physical environment of this coastal forest was likely mild and humid, perhaps even swampy.

The specific community where the fossils found in El Chango quarry lived was probably a mixed forest where *Sequoiadendron helicalancifolium* dominated the highest canopy strata, in the same way *Sequoiadendron giganteum* and *Sequoia sempervirens* do today in California. We can visualize *Microcachrys rhomboidea* as a shorter plant—perhaps even a shrub similar to the extant *Microcachrys tetragona*—associated with swampy areas, in the same way *Microcachrys novae-zelandiae* R.J.Carp., G.J.Jord., Mildenh. & D.E.Lee did in the Oligo-Miocene (Carpenter et al., 2011). *Dacrydium bifoliosus* probably occupied an intermediate stratum in the forest, and all three conifers coexisted with angiosperms, likely in the form of shrubs.

## CONCLUSIONS

Every study of new fossil assemblages offers a unique opportunity to visit the past. The co-occurrence of *Sequoiadendron helicalancifolium*, *Dacrydium bifoliosus*, and *Microcachrys rhomboidea* during the Cretaceous in the area that today is Chiapas, Mexico is a valuable insight from the southernmost North American Cretaceous ecosystems. The discovery of these fossils in El Chango adds to the fossil record of podocarpaceous and sequoioid conifers. In particular, *Microcachrys rhomboidea* is the oldest macrofossil of *Microcachrys*, filling in the expectation of finding this lineage in the Mesozoic based on the pollen-record. Finally, according to our interpretation, El Chango was a humid coastal conifer-dominated forest, supporting the hypothesis of a geographically and ecologically structured “rise of angiosperms” at the end of the Cretaceous, with conifers remaining dominant in coastal environments while angiosperms became dominant in freshwater-associated ecosystems. El Chango has proven to be a quarry with outstanding preservation quality. Given the relevance of the geographic and temporal location of this quarry, more paleontological and geological work here—in particular the study of the angiosperm assemblage, taphonomic studies, and fine-scale geology—will have a great impact in our understanding of the evolution of plant lineages and terrestrial ecosystems.

## Acknowledgments

The authors thank the members of the Paleobotany lab at UNAM and the Museo de Geología UNICACH for their help in the field. We especially thank the welcoming community of Pluma de Oro, in Ocozocuatla, Chiapas, who allowed us to establish our base campsite in their land and shared their water with us. We thank Enoch Ortiz for sharing his knowledge on the preparation of the fossil material, Laura Calvillo-Canadell for her support during the initial steps of this work, and Helga Ochoterena, David Gernandt, and Damon Little for their feedback through the initial stages of this project. We also thank Jenna T.B. Ekwealor, Tanner Frank, and Michael R. May for their feedback on the manuscript. Thank you to Aldo Dominguez for the illustration of El Chango in the Cretaceous. We appreciate the contribution of XXXX anonymous reviewers that improved the quality of the manuscript.

This research was supported by projects to SRSCF from Dirección General de Asuntos del Personal Académico (DGAPA: PAPIIT), UNAM, under Grant IN210416 and CONACYT under Grants 221129 and CF61501.

IGR was funded by CONACyT Master’s fellowship number 363004 to complete the program in Posgrado en Ciencias Biológicas at UNAM, and CJR was supported by the National Science Foundation (NSF) grant DEB-1754705. Part of the herbarium data collection at the NYBG was possible through CONACyT’s Beca Mixta program.

## Author Contributions

IGR curated the fossils, assembled the data matrices, conducted the phylogenetic analyses and wrote the first draft of the manuscript. SRSCF conceived the project, supported the fieldwork through his research grant, and reviewed the manuscript. CJR contributed to manuscript writing.

## Data availability statement

All the data and supplemental materials associated with this research are available in the GitHub repository: https://github.com/ixchelgzlzr/coniferas_el_chango And the Dryad repository **XXXXXX** (It will be updated once the manuscript is accepted).

