## Supplemental information for "Three new Cenomanian conifers from El Chango (Chiapas, Mexico) offer a snapshot of the geographic mosaic of the Mesozoic conifer decline"

### Supplemental Materials for: Three new Cenomanian conifers from the El Chango deposit (Chiapas, Mexico)

September 1, 2021

#### Contents

**S1 Extant terminals and Genbank references** **S2**

**S2 Morphological characters** **S3**

### S1 Extant terminals and Genbank references

| Family | Species | Author | rbcL | matK |
|---|---|---|---|---|
| Ginkgoaceae | *Ginkgo biloba* | L | AJ235804.1 | AF456370.1 |
| Taxaceae | *Cephalotaxus harringtonii* | Knight ex J.Forbes) K.Koch | JQ512524.1 | EF660650.1 |
| Taxaceae | *Torreya nucifera* | L. Siebold & Zucc. | JQ512624.1 | AB030137 |
| Taxaceae | *Taxus cuspidata* | Siebold & Zucc | EF660720.1 | AF228104.1 |
| Taxaceae | *Austrotaxus spicata* | Compton | AF456385.1 | AF456378.1 |
| Cupressaceae | *Cunninghamia lanceolata* | (Lamb.) Hook. | AY140260.1 | AB030125.1 |
| Cupressaceae | *Taiwania cryptomerioides* | Hayata | JF725919.1 | AB023999.1 |
| Cupressaceae | *Athrotaxis selaginoides* | D.Don | JF725938.1 | AB030130.1 |
| Cupressaceae | *Athrotaxis cupressoides* | D.Don | JF725921.1 | AB030131.1 |
| Cupressaceae | *Sequoia sempervirens* | (D.Don) Endl. | L25755.2 | HQ245909.1 |
| Cupressaceae | *Metasequoia glyptostroboides* | Hu & W.C.Cheng | AB917054.1 | HQ245903.1 |
| Cupressaceae | *Sequoiadendron giganteum* | (Lindl.) J.Buchholz | AY056580.1 | HQ245910.1 |
| Cupressaceae | *Cryptomeria japonica* | (Thunb. Ex L.f.) D.Don | L25751.2 | AB023984.1 |
| Cupressaceae | *Taxodium mucronatum* | C.Lawson | JF725913.1 | AB030119.1 |
| Cupressaceae | *Glyptostrobus pensilis* | (Staunton ex D.Don) K.Koch | JF725918.1 | AB030118.1 |
| Cupressaceae | *Actinostrobus arenarius* | (C.A.Gardner) Silba | JF725937.1 | JF725837.1 |
| Cupressaceae | *Actinostrobus pyramidalis* | Miq | JF725931.1 | HQ245874.1 |
| Cupressaceae | *Austrocedrus chilensis* | (D.Don) Pic.Serm. & Bizarri | EU161449.1 | HQ245876.1 |
| Cupressaceae | *Callitris macleayana* | (F.Muell) F.Muell | JF725933.1 | HQ245878.1 |
| Cupressaceae | *Callitris preissii* | Miq. | AY988230.1 | JF725840.1 |
| Cupressaceae | *Diselma archeri* | Hook.f. | l12572.2 | HQ245889.1 |
| Cupressaceae | *Fitzroya cupressoides* | (Molina) I.M.Johnst. | JF725916.1 | HQ245890.1 |
| Cupressaceae | *Fokienia hodginsii* | (Dunn) A.Henry& H.H. Thomas | EF053219.1 | HQ245891.1 |
| Cupressaceae | *Libocedrus plumosa* | (D.Don) Druce | JF725928.1 | AF152200.1 |
| Cupressaceae | *Libocedrus bidwillii* | Hook.f. | JF725927.1 | AF152202.1 |
| Cupressaceae | *Neocallitropsis pancheri* | (Carrière) de Laub. | AF127426.1 | AF152205.1 |
| Cupressaceae | *Papuacedrus papuana* | (F.Muell) H.L.Li | EU161451.1 | AF152206.1 |
| Cupressaceae | *Widdringtonia wallichii* | J.A.Marsh | AY140261.2 | JF725836.1 |
| Cupressaceae | *Tetraclinis articulata* | (Vahl) Mast. | L12576.2 | AF152213.1 |
| Cupressaceae | *Thujopsis dolabrata* | (L.f.)Siebold & Zucc. | JQ512621.1 | AF152217.1 |
| Cupressaceae | *Platycladus orientalis* | (L.) Franco | JQ512599.1 | AF152208.1 |
| Cupressaceae | *Thuja occidentalis* | L. | JQ512620.1 | AF152214.1 |
| Cupressaceae | *Juniperus drupacea* | Labill. | HM024301.1 | HM024023.1 |
| Cupressaceae | *Juniperus communis* | L. | HM024295.1 | HM024017.1 |
| Cupressaceae | *Juniperus rigida* | Siebold & Zucc. | JQ512551.1 | AB030136.1 |
| Cupressaceae | *Juniperus recurva* | Buch.-Ham. ex D.Don | HM024328.1 | HM024050.1 |
| Cupressaceae | *Juniperus squamata* | Buch.-Ham. ex D.Don | HM024339.1 | HM024061.1 |
| Cupressaceae | *Juniperus flaccida* | Schltdl. | HM024304.1 | HM024026.1 |
| Cupressaceae | *Juniperus monosperma* | (Engelm.) Sarg. | HM024315.1 | HM024037.1 |
| Cupressaceae | *Junipeus californica* | Carrière | HM024291.1 | HM024013.1 |
| Cupressaceae | *Juniperus virginiana* | L. | AF119182.1 | JQ512432.1 |
| Cupressaceae | *Juniperus bermudiana* | L. | HM024289.1 | HM024011.1 |
| Cupressaceae | *Juniperus phoenicea* | L. | HM024320.1 | HM024042.1 |
| Cupressaceae | *Cupressus funebris* | Endl. | AY988245.1 | HQ245888.1 |
| Cupressaceae | *Cupressus sempervirens* | L. | HM024278.1 | AF152187.1 |
| Cupressaceae | *Callitropsis nootkatensis* | (D.Don) Oerst. Ex D.P.Little | HM024268.1 | FJ475239.1 |
| Cupressaceae | *Hesperocyparis macnabiana* | (A.Murray) Bartel | AY380890.1 | AY380848.1 |
| Cupressaceae | *Hesperocyparis forbesii* | (Jeps.) Bartel | HM024281.1 | AY988343.1 |
| Cupressaceae | *Hesperocyparis sargentii* | (Jeps.) Bartel | AY988254.1 | AY9497215.1 |
| Cupressaceae | *Hesperocyparis macrocarpa* | (Hartw.) Bartel | AY380891.1 | AY380849.1 |
| Cupressaceae | *Calocedrus macrolepis* | Kurz | EF053220.1 | AF152179.1 |
| Cupressaceae | *Chamaecyparis pisisfera* | (Siebold & Zucc.) Endl. | HM024274.1 | HM023986.1 |
| Cupressaceae | *Chamaecyparis lawsoniana* | (A.Murray bis) Parl. | HM024272.1 | HM023984.1 |
| Cupressaceae | *Chamaecyparis obtusa* | (Siebold & Zucc.) Endl. | JQ512527.1 | AB030133.1 |
| Sciadopytiaceae | *Sciadopitys verticillata* | (Thunb.) Siebold & Zucc. | L25753.2 | AB023994.1 |
| Podocarpaceae | *Microcachrys tetragona* | (Hook.) Hook.f. | HM593611.1 | HM593713.1 |
| Podocarpaceae | *Dacrycarpus dacrydioides* | (A.Rich.) de Laub. | AF249597.1 | HM593702.1 |
| Podocarpaceae | *Halocarpus bidwillii* | (Hook.f. ex Kirk) C.J.Quinn | AF249638.1 | KF713609.1 |
| Podocarpaceae | *Lagarostrobos franklinii* | (Hook.f.) Quinn | AF249641.1 | HM593710.1 |
| Podocarpaceae | *Pherosphaera fitzgeraldii* | (F.Muell.) Hook.f. | AF249646.1 | HM593719.1 |
| Podocarpaceae | *Lepidothamnus laxifolius* | (Hook.f.) Quinn | HM593610.1 | AF457114.1 |
| Podocarpaceae | *Dacrydium cupressinum* | Corner | AF249634.1 | AF457112.1 |
| Podocarpaceae | *Dacrydium lycopodioides* | Brongn. & Gris | HM593596.1 | HM593695.1 |
| Podocarpaceae | *Podocarpus elongatus* | (Aiton) L´Hér. ex Pers | HM593643.1 | HM593746.1 |
| Podocarpaceae | *Retrophyllum comptonii* | (J.Buchholz) C.N.Page | AF249660.1 | HM593785.1 |
| Pinaceae | *Picea mariana* | (Mill.) Britton, Sterns & Poggenb. | EF440585.1 | EF440507.1 |
| Pinaceae | *Pinus taeda* | L. | AF119177.1 | AB080928.1 |
| Pinaceae | *Abies bracteata* | (D.Don) Poit. | AF456380.1 | AB080928.1 |
| Pinaceae | *Keteleeria davidiana* | (C.E.Bertrand) | EU269031.1 | AB161020.1 |
| Araucariaceae | *Araucaria araucana* | (Molina) K.Koch | AF249664.1 | AF456373.1 |
| Araucariaceae | *Araucaria hererophylla* | (Salisb.) Franco | U96462.1 | AF456374.1 |
| Araucariaceae | *Agathis australis* | (D.Don) Lindl. | JN627328.1 | JN627305.1 |

Table S1: Species of conifers used in the phylogenetic analysis. *Ginkgo biloba* was used as outgroup. The Genbank accession numbers are indicated for the two markers used

#### S2 Morphological characters

1. Phyllotaxy of first-order branches: (0) whorled to pseudowhorled, (1) helical, (2) irregular.
2. Last-order branch arrangement: (0) mostly tridimensional (1) mostly pinnate.
3. Type of mature leaves: (0) scale-like, (1) awl-like, (2) needle-like, (3) extended blade, (4)linear.
4. Leaves decurrence: (0) non-decurrent, (1) base decurrent and tip free, (2) totally decurrent
5. Presence of short branches: (0) absent, (1) present.
6. Scale-like leaves dimorphism (facial vs lateral): (0) absent, (1) present.
7. Node length uniformity: (0) regularly spaced, (1) irregularly spaced.
8. Presence of two different types of leaves in adult foliage: (0) absent, (1) present.
9. Projection in the tip of scalelike leaves: (0) present, (1) absent.
10. Phyllotaxy of mature predominant leaves; (0) helical, (1) opposite and decusate, (2) whorls of three leaves, (3) whorls of more than three leaves, (4) distichous.
11. Mature leaves in transversal section shape: (0) rhomboidal to tetragonal, (1) thin and broad.
12. Mature leaf apex form: (0) acute, (1) obtuse.
13. Mature leaf apex curvature: (0) straight, (1) curved.
14. Leaf curvature: (0) curved inwards, (1) curved outwards, (2) straight.
15. Mature leaf venation: (0) single middle vein, (1) two visible veins, (2) multiple parallel veins, (3) multiple reticular veins, (4) without visible veins.
16. Midvein from external view: (0) conspicuous, (1) inconspicuous.
17. Mature leaf-brach insertion: (0) sessile, (1) reduced base, (2) petiolate.
18. Mature leaf margin: (0) entire, (1) serrate, (2) minutely denticulate.
19. Resin gland presence: (0) absent, (1) present.
20. Resin gland type: (0) round and not subdermal, (1) oblonge extending along the leaf and subdermal.

21. Stomata disposition on mature leaves: (0) amphistomatic, (1) hypostomatic, (2) epistomatic.
22. Stomata organization: (0) scattered, (1) in one strip, (2) in two parallel strips, (3) in parallel rows of stomata.
23. Stomata shape: (0) round, (1) ovate.
24. Epidermal cells shape: (0) rectangular, (1) ovate, (2) irregular.
25. Epidermal cells organization: (0) in rows, (1) dispersed.
26. Florin rings: (0) present, (1) absent.
27. Frequent number of subsidiary cells: (0) 4-5, (1) more than 5, (2) fewer than 4.
28. Ovuliferous cone position: (0) terminal, (1) lateral.
29. Ovuliferous cone type: (0) simple, (1) compound.
30. Ovuliferous cone shape: (0) spherical/globose, (1) ellipsoidal/subglobose, (2) cylindrical, (3)irregular.
31. Ovuliferous cone texture at maturity: (0) mostly lignified (woody), (1) mostly coriaceous, (2) mostly fleshy.
32. Ovuliferous complex phyllotaxy: (0) helical, (1) decussate, (2) whorled.
33. Ovuliferous complex shape: (0) peltate, (1) non-peltate.
34. Sterile cone scales: (0) fewer than 4, (1) more than 4.
35. Collumela presence: (0) absent, (1) present.
36. Bract/scale fusion in the ovuliferous complex: (0) absent, (1) present.
37. Bract/scale degree of fusion: (0) totally fused, (1) relictual scale tip, (1) free scale up to one third of OC length, (2) free scale more than one third of OC length.
38. Ovuliferous complex ornamentation: (0) without ornamentation, (1) ornamentated.
39. Ovuliferous complex vascularization at origin: (0) independent trace for bract and scale, (1)unique trace for bract and scale.
40. Bract similar to vegetative leaves in ovulate cone base: (0) absent, (1) present.
41. Seed embedded in ovuliferous scale tissues: (0) absent, (1) present.
42. Ovule orientation at maturity: (0) perpendicular to ovulate cone axis, (1) parallel to ovulate cone axis, (2) oblique to ovulate cone axis.

43. Seed abscission: (0) absent, (1) present.
44. Integumentary seed wing: (0) abscent, (1) present.
45. Integumentary seed wing number: (0) one, (1) two, (2) three.
46. Integumentary seed wing symmetry: (0) asymmetric, (1) symmetric.
47. Seed position in ovuliferous complex: (0) adaxial, (1) terminal.
48. Epimatium: (0) absent, (1) present.
49. Cotyledons: (0) two, (1) three or more.
50. Pollen cone position: (0) lateral, (1) terminal.
51. Pollen cone aggregation: (0) absent, (1) present.
52. Pollen cone morphology: (0) spherical/globose, (1) ellipsoidal/subglobose, (2) cylindrical, (3)irregular.
53. Microsporophyll phyllotaxy: (0) decussate, (1) helical, (2) whorled.
54. Basal bracts on pollen cone: (0) absent, (1) present.
55. Life form: (0) shrub, (1) tree.
56. Foliage abcission: (0) evergreen, (1) deciduous.
57. Sex distribution in individuals: (0) monoecious, (1) dioecious.
